# Single-particle tracking of dynein identifies PP2A B55/SUR-6 as a cell cycle regulator of cortical force generation

**DOI:** 10.1101/2021.10.22.465443

**Authors:** Alan Edwards, John B. Linehan, Vincent Boudreau, Paul S. Maddox

## Abstract

Convergence and positioning of the pronuclei and mitotic spindle of many zygotes aids efficient division and is essential for early embryonic patterning. In the *C. elegans* zygote, interactions between microtubules and cortically anchored dynein are key to early development. However, how cortical microtubule pulling forces are controlled through the cell cycle is less well understood. We used single-molecule imaging and a windowed mean squared displacement analysis to uncover the behavior of dynein during cortical force generation, and provide a regulatory role for protein phosphatase PP2A-B55/SUR6 via NuMA-like protein LIN-5 in this process. Previous findings and our results suggest that PP2A regulates cortical microtubule pulling forces by increasing dynein binding and unbinding to the cortical force generation complex. Our data also suggests that cortical occupancy of dynein is abrogated to vary force generation. Our approach will be broadly applicable to classify the force generation behavior of single molecules in living organisms.

## Introduction

The minus-end directed microtubule motor protein dynein plays a key role in many cellular events. Dynein utilizes its ability to “walk” along microtubules in a polarized fashion to shuttle cargo, ranging in size from proteins or small complexes to entire organelles, in a directed manner. Dynein can also generate force on a microtubule by either bundling and cross-linking adjacent microtubules or by tethering to a membrane and pulling on microtubules (Reviewed in Roberts *et al*., 2013).

During cell division, dynein is essential for the correct establishment and maintenance of the mitotic spindle (Merdes *et al*., 1996, Inoué *et al*., 1998). Dynein tethered to the cortex pulls on microtubules emanating from centrosomes, allowing for dynamic spindle placement (Laan *et al*., 2012, Gusnowski and Srayko 2011). Dynein also has a critical role in kinetochore regulation, where it can inactivate the spindle assembly checkpoint by trafficking checkpoint-associated proteins away from the kinetochore-microtubule attachment (Howell *et al*., 2001, Tanenbaum *et al*., 2013).

In the *C. elegans* zygote, dynein is responsible for pronuclear and centrosomal migration and positioning. Dynein is unevenly distributed at the cortex throughout different stages of the cell cycle, providing the polarity-induced force imbalance that drives asymmetric cell division. Cortical dynein binds and pulls centrosomal microtubules, aiding centrosome positioning. In addition, dynein tethered to the nuclear envelope pulls on microtubules to drive the separation of centrosomes and the pronuclei to migrate and fuse, triggering nuclear envelope breakdown (De Simone *et al*., 2016, De Simone and Gönczy 2017).

The function and regulation of dynein within the context of the cell cycle have been well characterized through a variety of different methods. Single molecule assays have probed the motoring capacity of dynein and characterized its motoring capacity and activation *in vitro*. The regulation of dynein motoring has been determined by investigating the relationship between dynein’s capacity for processivity and the presence of co-factors such as dynactin and LIS-1 (Desantis *et al*., 2017, Huang *et al*., 2012, McKenney *et al*., 2014, Gennerich *et al*., 2007, reviewed in Roberts *et al*., 2013, reviewed in Reck-Peterson *et al*., 2018). However, it is unclear how dynein behavior during force generation events at the single-molecule scale in live cells compares to *in vitro* studies.

*In vivo* studies focused on the cell cycle have defined the components of the force generation complex tethering dynein to the nuclear envelope and cell cortex (Malone *et al*., 2003, Crisp *et al*., 2005, Okumura *et al*., 2018). These studies used a combination of fluorescent labeling, genetic manipulations, biophysical and biochemical approaches to identify microtubule associated proteins and dynein binding proteins that tether dynein and enable it to bind to microtubules. Laser ablation of microtubules have approximated the amount of force dynein exerts on microtubules by measuring changes in the speed of centrosome or chromosome movement (Elting *et al*., 2014, Fielmich *et al*., 2018). Fluorescence correlation spectroscopy and a “tube assay” that measured force generation through cytoplasmic membrane invagination showed that the asymmetric binding rate of dynein to the cortex is responsible for the polarity-induced force imbalance seen in the *C. elegans* zygote (Redemann *et al*., 2010, Rodriguez-Garcia *et al*., 2018).

In addition, mitotic kinases and phosphatases have been shown to regulate dynein polarization and recruitment to the cortex throughout stages in the cell cycle. In HeLa cells, Cyclin-dependent kinase 1 (CDK-1) negatively regulates cortical dynein accumulation through the phosphorylation of the dynein cortical anchor NuMA, and the protein phosphatase PP2CA counteracts the phosphorylation of NuMA by CDK-1 (Kotak *et al*., 2013). In the *C. elegans* two-cell embryo (Ab and P1), Polo-like kinase 1 (PLK-1) has been shown to negatively regulate the polarization and recruitment of dynein to the cortex by decreasing cortical accumulation of the dynein cortical anchor, NuMA-like protein LIN-5 (Bondaz *et al*., 2019). In the *C. elegans* zygote, protein phosphatase 2A (PP2A-B55/SUR-6, hereafter referred to as PP2A) is required for proper centrosome and pronuclear positioning, as well as proper centrosome breakdown in anaphase (Boudreau *et al*., 2019, Enos *et al*., 2018). However, it is unclear how PP2A interacts with dynein to regulate cortical force generation.

To investigate these questions, we developed a single-particle tracking analysis tool by combining live-cell, high spatiotemporal resolution microscopy and single particle tracking with a windowed mean square displacement (wMSD) analysis. Using the *C. elegans* zygote, we utilized a dynein heavy chain-tagged strain (eGFP::DHC-1) (Schmidt *et al*., 2017) to observe dynein near or at the cortex during pronuclear migration and anaphase with Total Internal Reflection Fluorescence (TIRF) microscopy (Axelrod 1981). Analyzing cortical dynein trajectories using a wMSD analysis enabled us to discern the activity of dynein at different points in time throughout its trajectory path length. Thus, we quantified the amount of time each individual dynein exhibited distinct behaviors. We measured the aggregate behavior of dynein force generation under genetic manipulation of specific regulators, and characterized dynein diffusion behavior in the anterior and posterior regions of the polarizing zygote. We depleted the dynein motor complex directly (Dynein light-chain 1 (DLC-1), regulators of dynein motoring capacity including dynactin (DNC-1) and LIS-1, microtubules via TBA-2, cortically through its anchor LIN-5, and the catalytic subunit of the master mitotic phosphatase PP2A (SUR-6), in order to better understand dynein behavioral regulation at the cortex. Through the implementation of a novel and robust strategy for investigating dynein behavior, we propose that PP2A regulates dynein force generation at the cortex. PP2A regulation may occur through an interaction with a NuMA-like protein, LIN-5. Our technique can also be extended to study the behavior of force generation in a variety of systems.

## Results

### A time-localized diffusion analysis enables quantification of cortical dynein particle behavior

To better understand how dynein behaves while generating force, we directly observed dynein’s movement through time. To assess the behavior of dynein at the cortex, we used a *C. elegans* strain with endogenously tagged dynein heavy chain (eGFP::DHC-1) (Schmidt *et al*., 2017). We then tracked particles through time and generated unique trajectories that were analyzed to generate a MSD curve given by the Lévy Flight model,

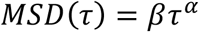

where *β* is the generalized diffusion constant, i.e. the medium surrounding a particle, *τ* is the time-lag with respect to an initial time point, and *α* describes the diffusion of the particle with respect to *τ*. Thus, *α* can be analyzed to determine the behavior of the particle through time with respect to *τ* (See Materials and Methods).

The *α* values resulting from an MSD analysis are thought of as belonging to one of three categories. When 0 < *α* < 1, *α* indicates constrained motion, *α* ≅ 1 indicates Brownian motion, and *α* > 1 indicates super diffusive motion (directed motion or applied force) (Metzler *et* al., 2014). It is also possible for the *α* value to be negative, as observed by Argyrakis *et al*. (2009).

We separately analyzed each category of diffusion between control and depletion. There was no observable difference in the time spent super diffusive, in brownian, or constrained diffusion categories. The negative diffusion modality is the only diffusive regime showing a difference between conditions. Given these results, we wanted to further probe dynein behavior associated with negative *α* values.

A negative *α* value is calculated when an object’s average displacement decreases as time-lag window increases. The object’s next position is likely to be closer to where it was at an earlier time (Figure 1 A, Supplemental Figure 1). Negative diffusion is also present in simulations of systems containing particles of different sizes. When the system is shifted from a higher temperature to a lower temperature (phase transition or condensate formation) the particles segregate by size, causing smaller particles to aggregate together. This aggregation of particles is the negative diffusion (Argyrakis *et al*.,2009).

**Figure 1.**
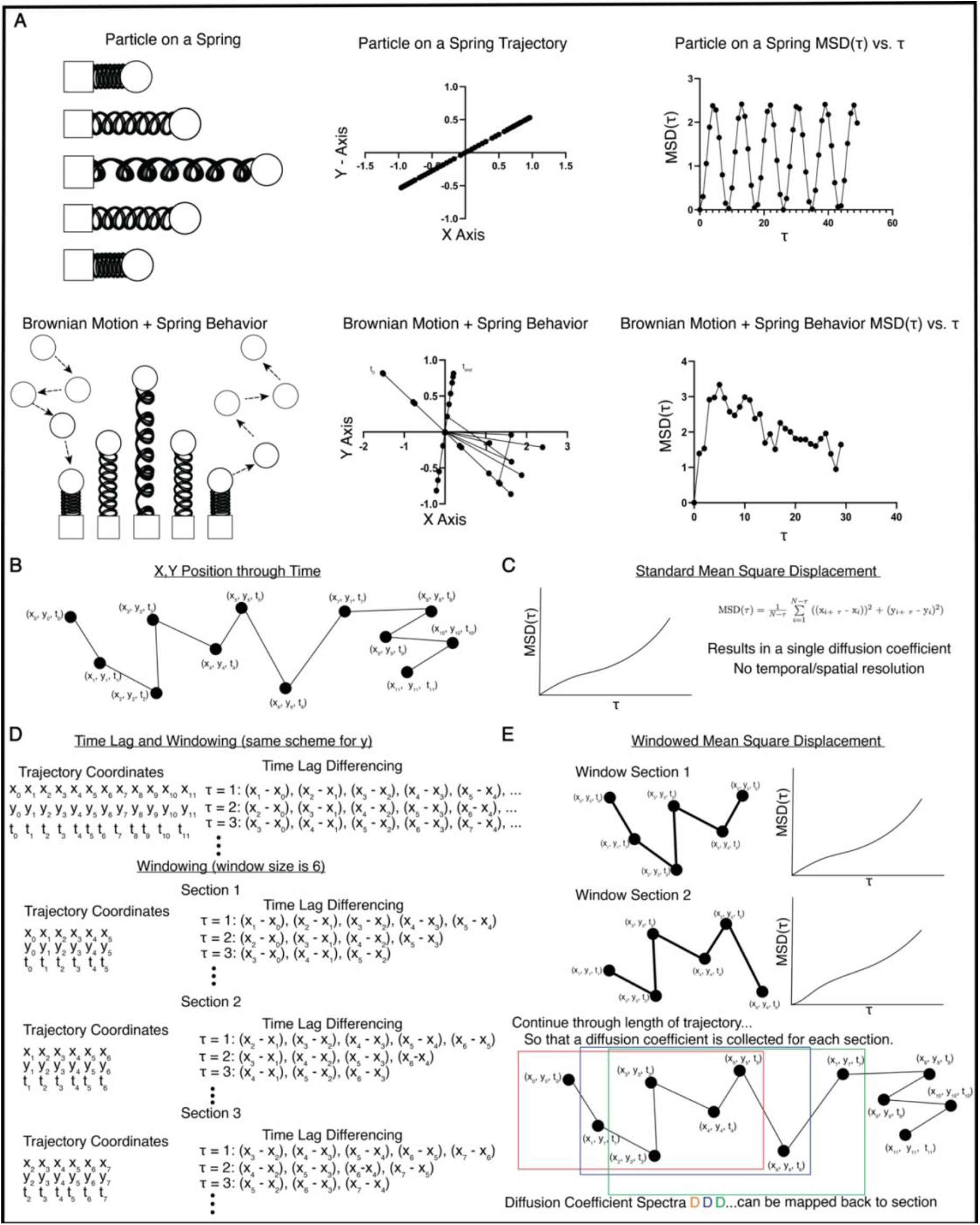
Schematic representation of windowed Mean Squared calculation analysis. A) Theoretical particle on a spring displaying oscillatory behavior (Top Panel). The MSD of the particle contains a regular interval of positive and negative slopes (Diffusion Coefficient). Theoretical particle on a spring with thermal fluctuations (brownian motion) (bottom panel). The MSD of the particle contains irregular intervals of positive and negative slope. B) Example trajectory in x,y position through time. C) Standard mean squared displacement (MSD) analysis is calculated using the equation in panel D, and results in a plot of the MSD versus time lag. D) Comparison of time lag using MSD analysis versus wMSD analysis. The MSD takes the time lag and applies it to all positions in the trajectory. The wMSD computes the time lag difference over a single section (n=6). E) The wMSD is the application of the MSD to a single section of the trajectory, from which a diffusion coefficient is calculated, providing a spatially resolved measure of motility.

For dynein to generate cortical force, dynein must be tethered to the cortex and to a microtubule, and dynein must also be active and persist in the state long enough to generate force. During force generation, we imagined dynein would be locally constrained, but would experience a “tug of war” of forces from its association with both the microtubule and cortex. These forces would result in a dynein particle that oscillates around a central point while dynein is generating force. Therefore, we reasoned that dynein behavior during force generation would be most accurately described by a negative *α* value.

However, we wanted to extend the sensitivity of this analysis, as dynein shows dynamic behavior throughout a single trajectory. Cortical dynein motility was quantified using a sliding window Mean Square Displacement (wMSD) (Korshidi *et al*., 2011). Rather than averaging the slope of MSD curve over an entire trajectory to generate a diffusion curve, the wMSD technique measures *α* at distinct intervals within a trajectory (Fig 1 B-E). The wMSD approach allowed us to interpret the behavior within a single dynein’s trajectory through the use of time localized diffusion coefficients. The magnitude and sign of *α* is a read out of distinct dynein behaviors such as times at which dynein is tethered to the cortex, dwelling at the cortex, or walking along a microtubule. The standard MSD generates a single *α* by averaging displacements over a number of time-lag windows, providing a quantification of the average behavior of dynein while localized near the cortex (Figure 1 B, C). By introducing a sliding window, spectra of *α* values are generated that describe dynein behavior as it follows through its trajectory (Figure 1 B, D, & E). The sliding window provides access to temporally distinct behaviors of the MSD analysis. We used this approach to calculate the amount of time a single dynein is tethered to the cortex (negative *α*), revealing changes in dynein behavior throughout its trajectory.

### Single-particle tracking of cortical dynein reveals different behaviors while localized to the cortex

To assess dynein behavior during cortical force generation, we utilized the single-cell *C. elegans* zygote, where cortical force generation across the anterior-posterior axis is asymmetric during different stages of the cell cycle (Grill *et al*., 2003, Labbé *et al*., 2004).

Beginning at the onset of centrosome separation, microtubule-based cortical force generation is accomplished through a complex consisting of G-alpha proteins (GOA1/2), LGN-like protein GPR1/2, NuMA-like protein LIN-5, and dynein (Fielmech *et al*., 2008, reviewed in Rose and Gönczy 2014). By combining high spatiotemporal resolution TIRF microscopy (∼37.4 frames/second, 0.06 μm/pixel) with a fluorescently tagged dynein heavy chain *C. elegans* strain (DHC-1::eGFP) we visualized dynein behavior near the coverslip (Figure 2 A). Using this method, we are able to track the dynamic movements of individual dyneins as they come in close proximity to the cortex (Figure 2 A, B, Movie 1).

**Figure 2.**
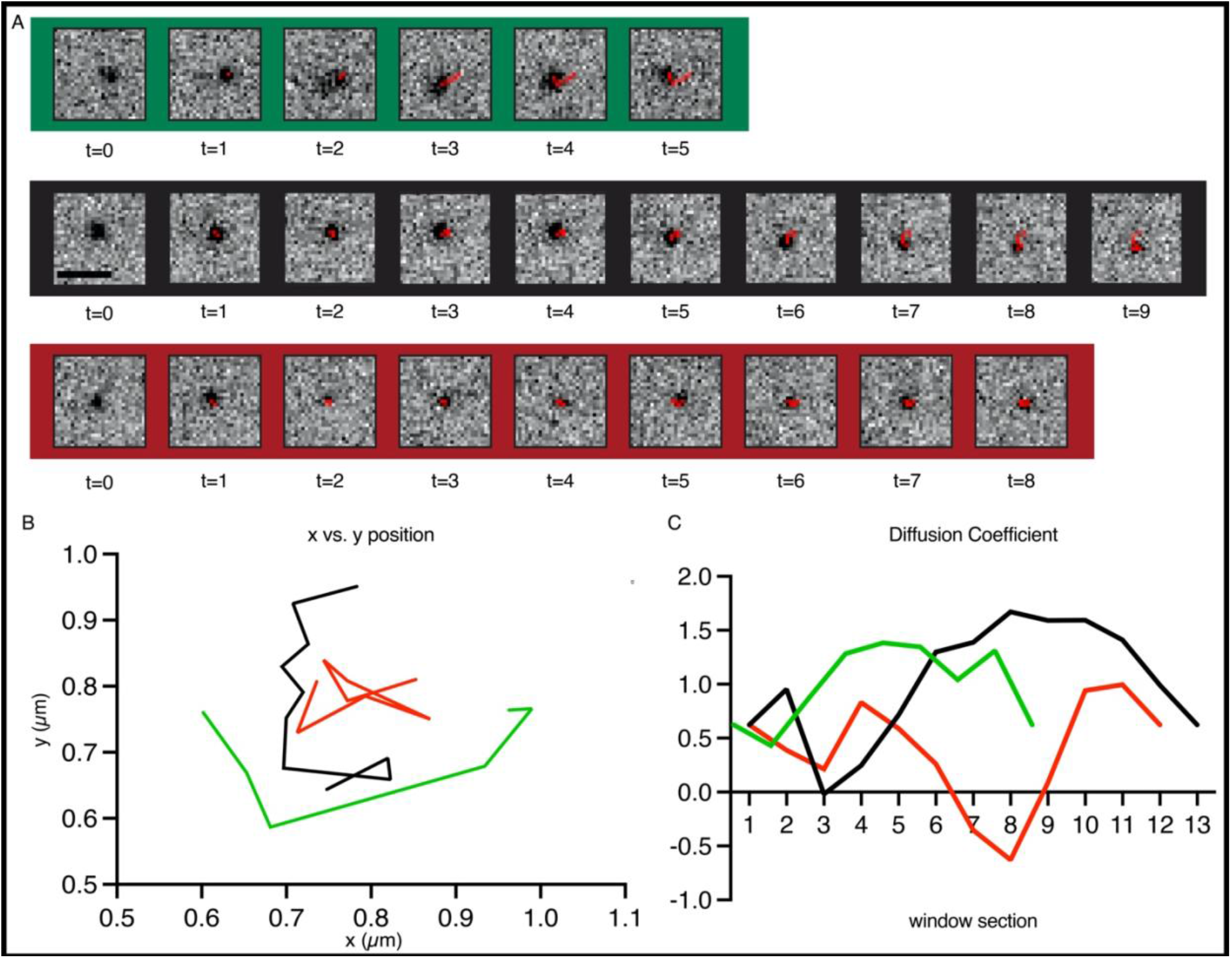
Analysis of wMSD from X,Y positioning. A) Three representative time-lapse tracking images of dynein (black dots) at the cortex using TIRF microscopy. B) Graphical recapitulation of x,y coordinates for the three representative tracks. C) Diffusion coefficients using a wMSD analysis extrapolated from the x,y positioning. Green = always positive diffusion, highest proportion of time spent with α> 1 (super-diffusive), Black = Diffusion from α<1 to α>1 (wide spectrum range of diffusion within the track), Red = Exhibits diffusion where α < 0 (negative diffusion). A single time point (t) = 0.027s. Scale bar = 1 µm.

### Single-particle tracking of asymmetric force generation in the *C. elegans* zygote suggests negative α within the diffusion spectra of dynein is the behavior responsible for cortical force generation

In *C*. elegans, the zygote is polarized and undergoes asymmetric cell division. Dynein-mediated cortical force generation is initially higher in the anterior during mitotic entry (Park and Rose 2008). If negative *α* indicates that dynein is tethered to the cortex and capable of binding to a microtubule, and thus, generating force, then negative *α* should be higher in the anterior during pronuclear migration. We didn’t observe any trends between conditions with other modes of *α* (data not shown).

To determine the efficacy of the wMSD diffusion spectra as a means of measuring cortical dynein force generation in *C. elegans*, we took advantage of the naturally-occurring asymmetry of cortical force generation in the zygote. Cortical dynein trajectories were measured from *C. elegans* zygotes in the single-cell stage during pronuclear migration. Cortical force generation is roughly two times greater in the anterior side during pronuclear migration than in the posterior side of the zygote (De Simone *et al*., 2017, De Simone & Gonczy, 2018, Rodriguez-Garcia *et al*., 2018). Thus, we expected to see that cortical dynein spends more time tethered to the anterior cortex than to the posterior cortex during pronuclear migration. Because α values resulting from the wMSD analysis were well localized in time, in that they can be mapped back to discrete sections of the trajectory, the time each dynein trajectory spends exhibiting negative *α* was calculated. The total time spent negative in the anterior and posterior of each zygote was calculated by summing up the time each dynein spends producing negative *α*. The average of the total time spent negative in anterior and posterior ends was calculated over each replicate. In zygotes, the average amount of time dynein is tethered to the cortex (having negative *α*) was 12.095 seconds in the anterior half of the cell (see materials and methods). In the posterior end the average time spent negative was 5.819 seconds (Figure 3 A, Figure S2A for statistical significance). On average, cortical dynein in the anterior end of the *C. elegans* zygote during pronuclear migration spends about two-fold more time tethered to the cortex than the posterior end, consistent with previous findings (De Simone *et al*., 2017, De Simone & Gonczy, 2018, Rodriguez-Garcia *et al*., 2018). These data suggest the average time spent negative calculated from the α spectra resulting from a wMSD analysis is an effective proxy for measuring cortical dynein force generation. To evaluate the significance of changes in average time spent negative that we observed between conditions, we performed a bootstrap statistical analysis, and compared bootstrapped values to measured values using a two-sample Kolmogorov-Smirnov test (Figure S2A for statistical significance, see materials and methods). Values for the average time spent negative were significantly different than what could be expected due to chance for all groups with the following exceptions: Anterior *dnc-1* (RNAi) and *sur-6* (RNAi), posterior *lis-1* (RNAi).

**Figure 3.**
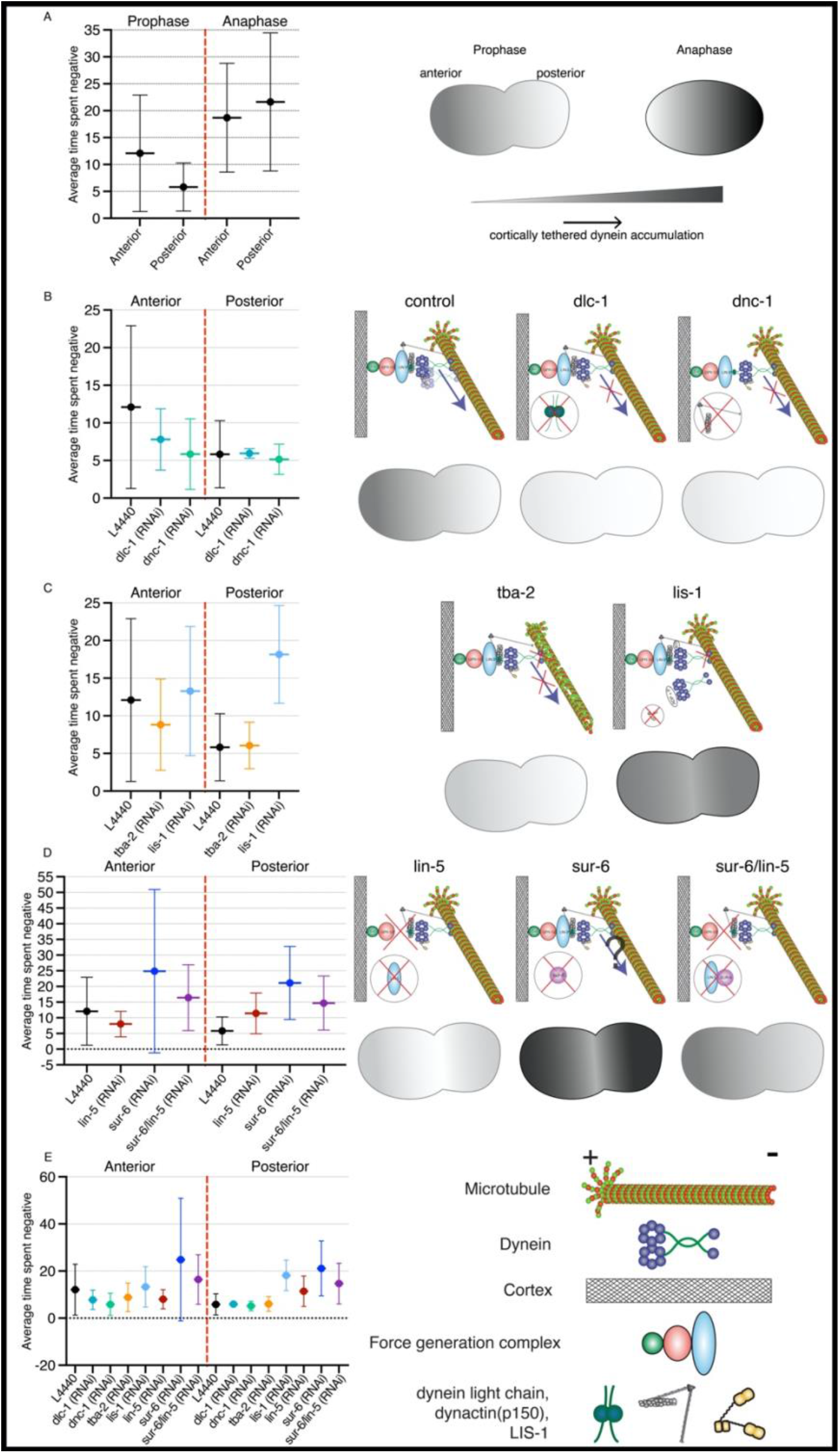
Average time spent in negative α (ATSN). A) ATSN is calculated in anterior and posterior segments of the zygote during pronuclear migration and anaphase. B) ATSN is calculated for depletions directly affecting dynein strucutre (DLC-1) and motoring (DNC-1) in the anterior and posterior segments of the zygote during pronuclear migration. C) ATSN is calculated for depletions directly affecting microtubule binding (TBA-2, LIS-1). D) ATSN is calculated for depletions affecting cortical binding (LIN-5), the catalytic subunit of PP2A (SUR-6), and the combined depletion of LIN-5 and SUR-6. E) Control and all depletions shown together. The amount of time dynein spends is recapitulated by a gradient for each condition. A schematic for each depletion is also provided for clarity.

To further test our approximation of force generation via negative *α*, we sought to measure dynein force generation during a different time in the cell cycle. In the *C. elegans* zygote, the asymmetry in force generation transitions from being greater in the anterior prior to pronuclear meeting, to being greater in the posterior during the metaphase to anaphase transition and through anaphase, resulting in a smaller P_1_ cell that is functionally and molecularly distinct from the Ab cell (Park and Rose 2008). This asymmetry gives rise to the differentiation of cell fate during development. Therefore, we measured dynein wMSD during anaphase, where force generation is higher in the posterior (Grill *et al*., 2003, Grill and Hyman 2005, Park and Rose 2008). We found that the force asymmetry measured by the average time spent negative in the anterior and posterior was 18.692 and 21.621 seconds, respectively (Figure 3A, Figure S2A for statistical significance). This result is consistent with findings based off of fluorescence intensity cortical localization of the force generating complexes GPR-1/2 and LIN-5 during anaphase in the single-cell zygote (Park and Rose 2008), confirming the validity of our approximation of force generation using average time spent negative.

### The loss of a dynein subunit or its motoring adaptor, dynactin, destabilizes dynein cortical force generation behavior

Having established negative α as a proxy indicator of dynein cortical force generation, we proceeded to investigate the effects of targeted depletions of dynein motor function. Depletion of subunits associated with dynein motoring, and thus microtubule binding, is expected to reduce the average total time spent with negative α. Dynein heavy chain 1 (*dhc-1*) is required for processivity. We chose to use a fluorescently labeled dynein heavy chain (DHC-1::EGFP), assuming that depletion of this subunit would result in complete loss of motoring. Instead, we depleted the dynein light chain subunit (DLC-1), with the intention of decreasing dynein function without abolishing force generation. The dynein subunit was depleted using *dlc-1*(RNAi) and fed to worms for 24 hours. Dynein trajectories collected after dlc-1 depletion had an average time spent negative in the anterior zygote of 7.794 seconds. The average time spent negative in the posterior of the zygote was 5.940 seconds (Figure 3B, Figure S2B for statistical significance). Comparison to L4440 control data, where anterior average time spent negative was 12.095 seconds, and posterior average time spent negative was 5.819 seconds, indicates that dynein force generation is more severely impeded by this DLC-1 depletion in the anterior end of the zygote. We hypothesize that the posterior end is consistent between the groups due to the limited availability of cortical force anchoring proteins such as GPR1/2 and LIN-5 in the posterior. (Rodriguez-Garcia *et al*., 2018, Park and Rose 2008). Thus, the RNAi depletion has the ability to interfere more with the anterior end since there is a larger number of dynein molecules generating force at that end of the zygote. Reduction in average time spent negative in *dlc-1*(RNAi) depleted zygotes in the anterior in comparison to the L4440 control data supports previous findings (Figure S2B for statistical significance, O’Rourke *et al*., 2007, Rodriguez-Garcia *et al*., 2018).

Next, we depleted dynactin via *dnc-1* (RNAi). DNC-1 is required for the processivity of dynein motility (Reck-Peterson *et al*., 2006). Upon 24 hour depletion of DNC-1, we observed the average time spent in negative α for the anterior and posterior as 5.874 seconds and 5.1660, respectively, a greater than two-fold reduction in cortical tethering in the anterior as compared to control. In *dnc-1* (RNAi) embryos, our statistical analysis showed that the regulation of dynein in the anterior is impeded in the absence of dynactin, resulting in dynein spending less time at the cortex, and only interacting with the cortex stochastically (Figure 3B, Figure S2B for statistical significance).

Finally, we depleted the microtubule/dynein-associated protein LIS-1, which has been shown *in vitro* to increase dynein binding affinity to a microtubule; therefore, in the absence of LIS-1, dynein exhibits more rapid unbinding rates (Huang 2012). However, LIS-1 is not required for dynein localization to the cortex *in vivo* (Cockell *et al*., 2004). Finally, *lis-1* (RNAi) zygotes display centrosome separation defects that phenocopy a decrease in motoring activity (Boudreau *et al*., 2019). The average time spent negative is notably increased in *lis-1* (RNAi), 13.288 seconds in the anterior and 18.160 seconds in posterior. Our statistical analysis indicated aberration of dynein regulation in the posterior (Figure 3C, Figure S2C for statistical significance). Depletion of *lis-1* results in increased dynein cortical tethering. Depletion of both *dlc-1* and *dnc-1* results in decreased dynein negative diffusive activity. Taken together, we show that depleting factors involved in dynein motoring abrogates the average amount of time dynein spends displaying negative α at the cortex and impedes dynein regulation.

### Disrupting microtubule polymerization decreases dynein force generation behavior

We next wanted to observe cortical dynein behavior in the presence of disrupted microtubule polymerization, where cortical force generation should be reduced. To do this, we depleted tubulin alpha chain-2 (TBA-2), which is a partially redundant isotype of the tubulin alpha chains, but is known to effect meiotic and mitotic spindle formation and microtubule dynamics in the early zygote (Lu and Mains, 2005; Honda *et al*., 2017). 24-hour depletion of *tba-2* (RNAi) reduced the zygote’s ability to establish polarity or were nonviable (data not shown). Thus, we depleted *tba-2* (RNAi) from zygotes for only 16 hours, which was penetrant, but non-lethal. As expected, we observed a decrease in the average time spent negative in the anterior (8.820 seconds), while there was a slight increase in the posterior (6.057 seconds) in comparison to control (Figure 3C, Figure S2C for statistical significance). This indicates that disrupting microtubule dynamics decreases dynein’s ability to generate cortical force in the anterior.

### Depletion of LIN-5 decreases the proportion of dynein cortical force generation behavior in the anterior

Dynein is anchored to the cortical force generation complex through NuMA like protein LIN-5, and it has been directly linked to dynein cortical accumulation and force generation (Park and Rose 2008, Fielmich *et al*., 2018). In order to assess how dynein behaves when its ability to bind to the force generation complex is reduced, we measured dynein diffusivity in *lin-5* (RNAi) zygotes. Again, we observed a strong reduction in the average time spent negative (tethered to the cortex) in the anterior, 8.017 seconds. We found an increase in the average time spent negative (tethered to the cortex) in the posterior, 11.43 seconds (Figure 3D, Figure S2D for statistical significance). Given that cortical force generation is roughly 1.5 times greater in the anterior during mitotic entry in the *C. elegans* zygote, we attribute this transition in force imbalance to the distribution of LIN-5 in a *lin-5* (*RNAi*) depletion scenario (Rodriguez - Garcia *et al*., 2018). We propose that actomyosin-based cortical flows are unable to redistribute the reduced pool of LIN-5 remaining from the depletion, resulting in increased average time spent negative in the posterior (discussed further in Discussion.)

### Loss of SUR-6 disrupts normal cortical dynein behavior

The master mitotic regulator and main counteracting phosphatase of Cyclin-Dependent Kinase 1 (CDK-1) is Protein Phosphatase 2A (PP2A). PP2A is a tripartite phosphatase consisting of a scaffold, catalytic (B55), and regulatory (SUR-6) subunit. PP2A is known to display high activity during onset of mitosis and at anaphase onset. It has been shown that its regulatory subunit SUR-6 is necessary for proper centrosome breakdown during anaphase and centrosome maturation in interphase (Enos *et al*., 2018). However, the mechanistic link between dynein and the force generation complex with SUR-6 is unknown. Hence, we wanted to assess cortical dynein behavior in *sur-6* (RNAi) zygotes. We found an increase in average time spent negative in the anterior (26.20 seconds) and posterior (24.86 seconds) (Figure 3D, Figure S2D for statistical significance). Our statistical analysis also identified a loss of dynein regulation at the anterior cortex in *sur-6* (RNAi) embryos (Figure S2D).

Given that *sur-6* (RNAi) decreases cortical microtubule polymerization velocities, our data suggests that dynein is able to bind to the force generation complex, but dynein’s ability to exert force on a microtubule is perturbed (Boudreau *et al*., 2019). This result leads to two potential ways by which SUR-6 regulates cortical dynein. SUR-6 could regulate dynein similarly to LIS-1, where a difference in the binding and unbinding rates of dynein to microtubules causes slippage and dynein fails to generate force. Given that SUR-6 does not directly regulate dynein motoring capacity at the nuclear envelope, we find this possibility unlikely (Boudreau *et al*., 2019). Alternatively, SUR-6 may also regulate cortical dynein indirectly, such that dynein can interact with microtubules and the cortex, but is unable to produce force on a microtubule.

### Single-particle tracking and RNAi depletions suggest a mechanistic relationship between the force generation complex and PP2a through the dephosphorylation of LIN-5 via SUR-6

To resolve previous observations taken from cortical tracking of *sur-6* (RNAi) in EBP2:GFP and EGFP::DHC-1 zygotes, we set out to find if SUR-6 interacts with the component that anchors dynein to the force generation complex, LIN-5. CDK-1 has been shown to negatively regulate the cortical force generation complex by phosphorylating LIN-5 (Portegijs *et al*., 2016, Kotak *et al*., 2013). As CDK-1’s main counteracting phosphatase, we reasoned that PP2A-B55/SUR-6 might positively regulate the cortical force generation complex through the dephosphorylation of LIN-5. In order to test this hypothesis, we simultaneously depleted SUR-6 and LIN-5 in DHC1:EGFP zygotes. If the two proteins do not interact, we expect to find an additive relationship between the individual depletions and double depletions in the time spent in negative α. If they do interact, we expect the double depletion of LIN-5 and SUR-6 to dampen the effects seen by the *sur-6* (RNAi) zygotes alone. We find that the double depletion of SUR-6/LIN-5 substantially reduces the time spent in negative α compared to SUR-6 zygotes, partially rescuing the *sur-6* (RNAi) phenotype (average time spent negative_sur-6 A_= 24.861 seconds, average time spent negative_sur-6/lin-5 A_= 16.406; average time spent negative_sur-6 P_= 21.132 seconds average time spent negative_sur-6/lin-5 P_= 14.685 seconds) (Figure 3D, Figure S2D for statistical significance). This data suggests that SUR-6 regulates cortical force generation by antagonizing LIN-5.

### Intensity measurements auto correlate diffusion orthogonal to the cortex

The analysis so far has used the x/y position of dynein throughout its trajectory to determine if the molecule is in a position to generate force at the cortex (tethered to the cortex). Since cortical dynein is free to move in three dimensions, we needed a way to infer movement in the z-direction to better understand dynein activity at the cortex when generating force. The fluorescent intensity of the GFP marker is used as a way of inferring the z-position of cortical dynein. Given that TIRF microscopy creates an exponentially decaying excitation, we expect that fluorescence intensity can be used as a measure to approximate the z-position of dynein (i.e., where dynein is within the excitation gradient, dynein close to the cortex will have a greater fluorescence intensity than dynein further away). We postulated that since dynein is constrained by tethering to the cortex in (x,y), dynein is also constrained to a small range of motion in the z dimension (Figure 4A). Having collected trajectories of cortical dynein, we could then analyze the fluorescence intensity data using the same wMSD analysis.

**Figure 4.**
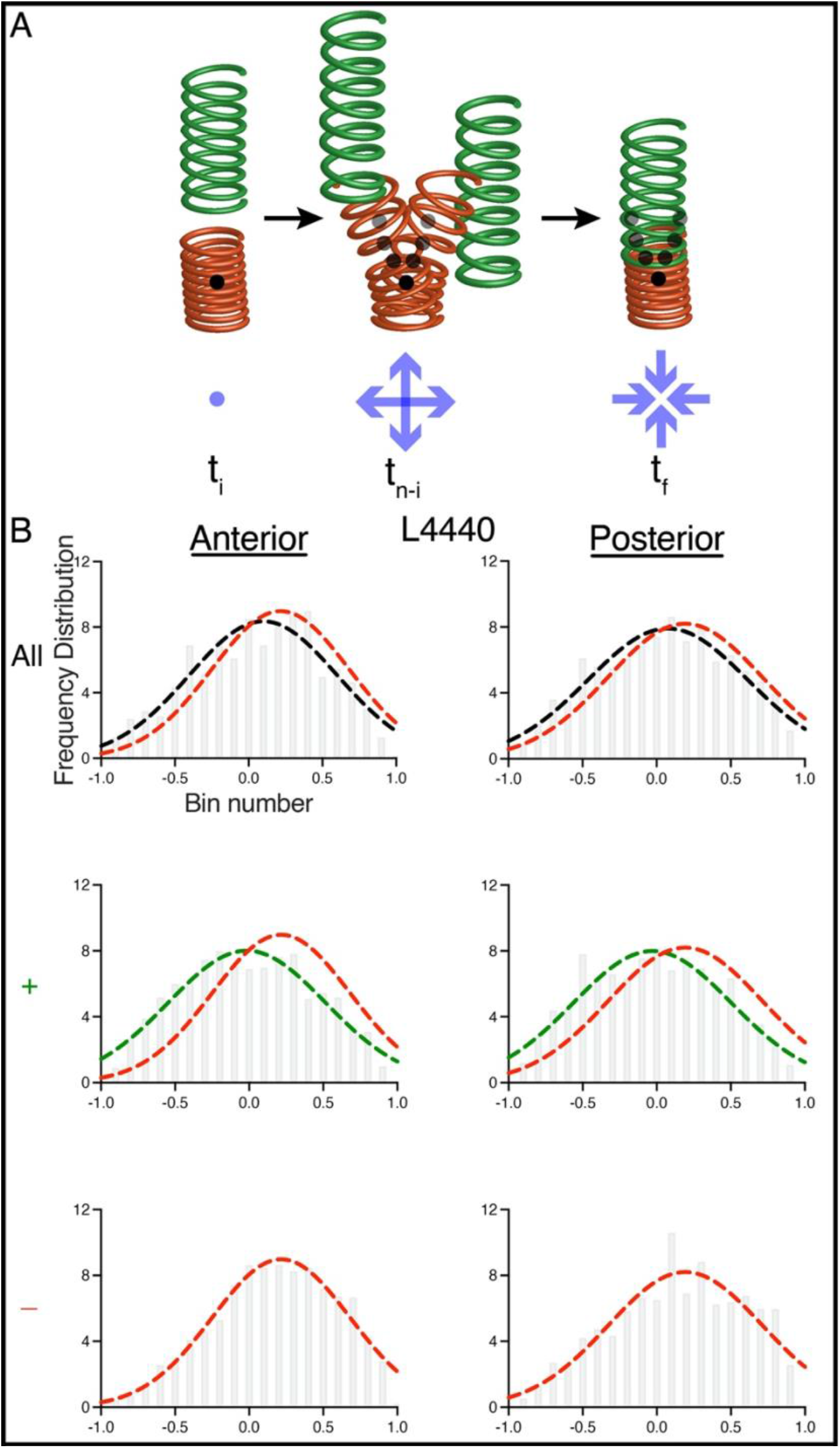
Cross Correlation of x,y diffusion with intensity (z) diffusion. A) Theoretical model of dynein force generation. An intial movement away from the intial point is coupled with movement back towards the initial point through time in x, y, and z planes. B) Histogram of cross correlation anaylsis for control (L4440). Comparison of all diffusion tracks shows no correlation (top, black). Positive diffusion in the x,y plane (middle green) reveals a negative correlation with positive expectation value. Negative diffusion (bottom, red) is in x,y and z dimensions. Gaussian distribution fits were generated for each histogram, and the negative diffusion curve is overlayed in red for comparison.

Extensive image processing was performed to acquire trajectories from the data that we could be confident had accurate (x, y) position information and were nearest to the cortex (materials and methods). A wMSD analysis was performed on the (x, y) positions and fluorescent intensity separately. We reasoned that treating both (x, y) and fluorescence intensity data with wMSD affords the ability to assess the correlation between the resulting α spectra from each data set.

For each dynein trajectory, we generated an α spectrum for its (x, y) position and fluorescence intensity data. We then calculated a correlation coefficient to determine if the two α spectra had a similar trend. We then made a histogram for each correlation coefficient from every trajectory. For simplicity, we only report the cross-correlation for the control (L4440). We did not observe any difference in the cross-correlation data amongst conditions (data not shown). When all the trajectories were included, we observed a left skewed distribution (Figure 4B, top panel). To determine the source of the skew in the total population histogram, we analyzed those tracks that did tether to the cortex, and tracks that never tethered to the cortex, separately. To determine the source of the skew, we generated a histogram of trajectories that only had positive α values (never tethered to the cortex) in (x, y) α spectra (Figure 4B, middle panel). These values are normally distributed about zero, thus the trajectories are not correlated in (x,y) and z. We then looked at the distribution of correlation coefficients that had negative α (tethered to the cortex) values in the (x, y) α spectra (Figure 4B, bottom panel). The histogram for the correlation coefficients of trajectories that did tether to the cortex (negative *α* in x, y plane) was a left-skewed distribution with the majority of values shifted to the right (positive). In the histogram of correlation coefficients for trajectories with negative *α* (tethered to the cortex) in the x,y plane, we saw that the expectation value of the distribution is positive. Thus for the population of dynein trajectories that did tether to the cortex, there was a correlation between movement in x,y and z. The trajectories that are tethered to the cortex are the source of the skewness in the total population histogram of correlation coefficients.

Thus, we expect that negative *α* values measured in (x,y) are a readout of force generation behavior, given the correlation of the (x,y) *α* spectra with the *α* spectra in z. Movement in (x,y) correlates with movement in z when dynein tethers to the cortex (negative *α*).

These results suggests that small oscillations of dynein in both the (x,y) and z accompany cortical-force generation, leading us to propose a simple spring-like tethering model for the behavior of cortically anchored dynein exerting force on a microtubule (Figure 6A). The implications of this model from a mechanistic standpoint will be discussed further below.

**Figure 5.**
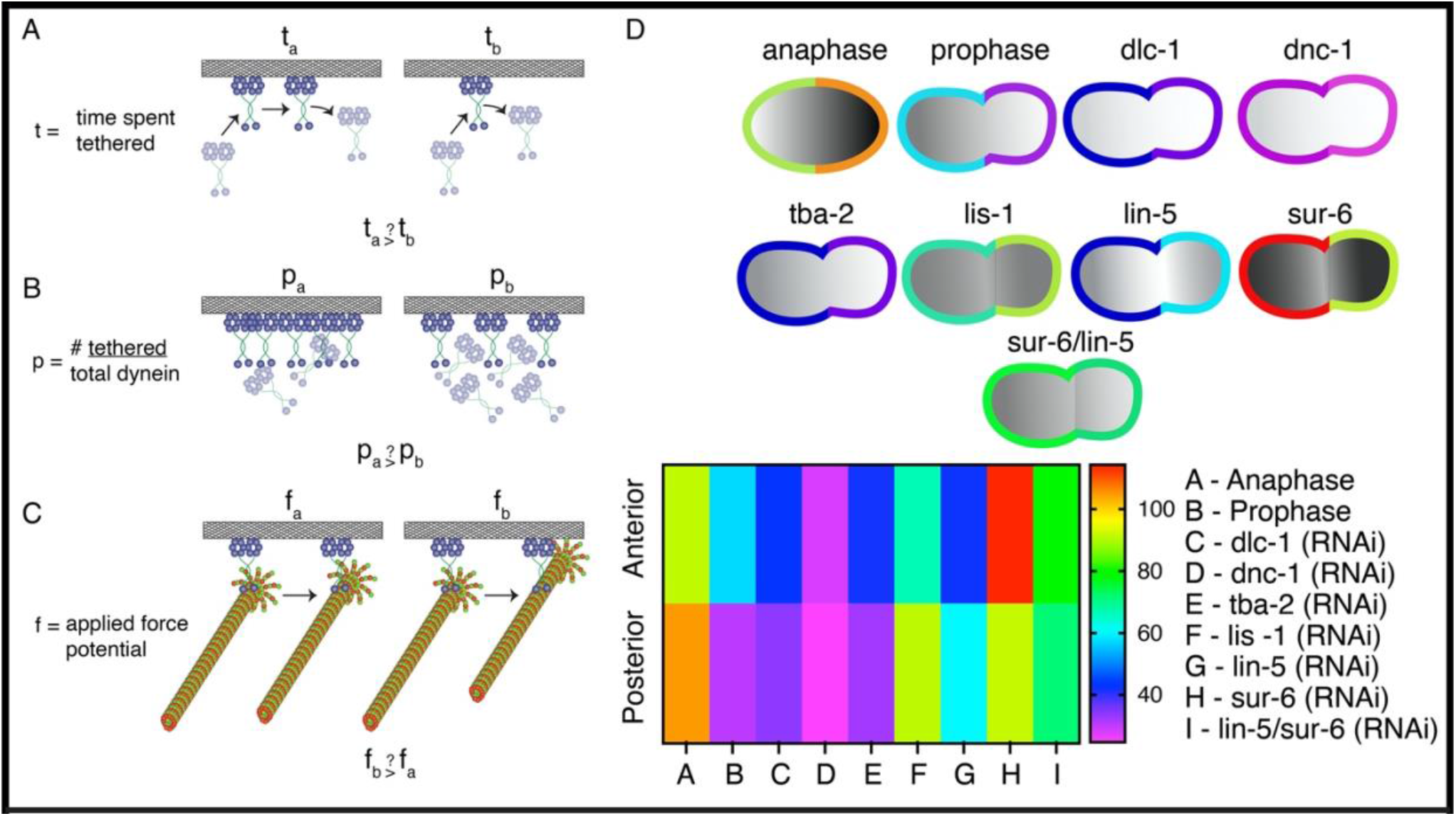
The number of dynein particles exhibiting tethering behavior is regulated during cortical dynein force generation. Variation in dynein cortical force generation is due to the: A) time individual dynein particles spend tethered to the cortex, B) Number of dynein particles tethered to the cortex, or C) force generating capacity (potential) of dynein in differnet regions. D) Heatmap of anterior and posterior values for number of dynein particles displaying negative α behavior at any point during individual trajectories.

**Figure 6.**
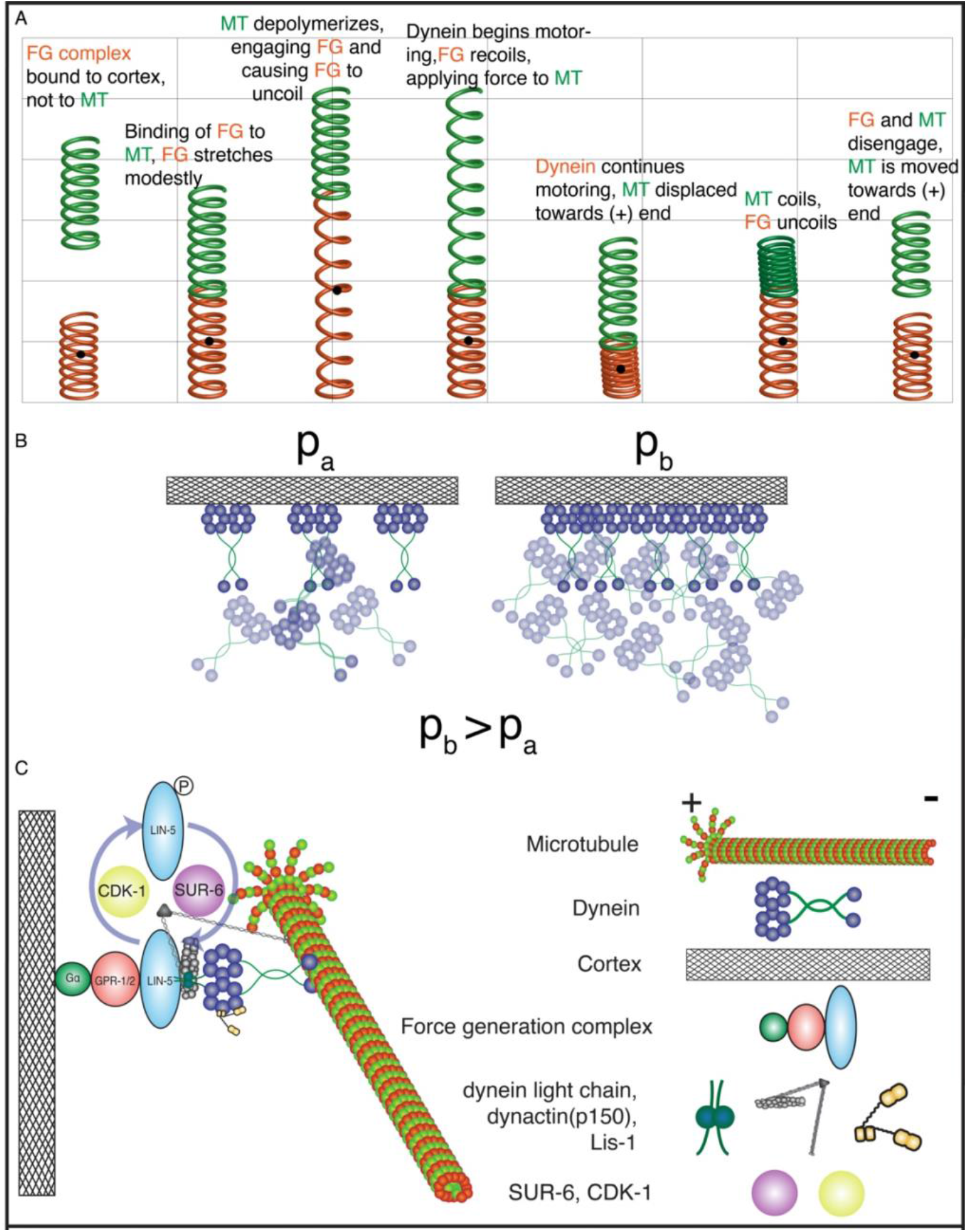
Model for Dynein force generation and regulation at the cortex. A) A simple spring-like model can explain the behavior of cortical dynein force generation. Force generation complex = FG. B) The number of dynein particles recruited to the cortex is positively correlated with the number of dynein particles engaging in force generation behavior. C) A putative mechanistic model of CDK-1 and PP2a (SUR-6) regulation of cortical dynein force generation via phosphorylation and dephosphorylation via LIN-5.

### The number of dynein particles exhibiting tethering behavior is regulated during cortical dynein force generation

We wanted to better understand why we observed variations in the average time dynein spent capable of force generation across conditions. We reasoned there could be at least four possibilities for these observations: 1) Dynein particles may exhibit different lengths of time displaying tethering behavior (Figure 5A), 2) The number of dynein particles in a population exhibiting tethering behavior varies (i.e. binding rate) (Figure 5B), 3) Dynein particles in different regions (i.e. anterior and posterior) may vary in force generation potential, such that a single dynein particle may produce a greater magnitude of force in the anterior versus the posterior (Figure 5C), and 4) dynein particles undergo a combination of influences arising from possibilities 1, 2, and/or 3. To identify a potential mechanism responsible for variations in dynein negative α spectra across conditions, we first measured the mean and median time individual dynein trajectories spent displaying negative α behavior, when any negative α behavior was observed within the trajectory. There were no distinguishable variations between conditions, with all conditions centered (median) around 0.189s (Supplementary Table S1). We then wanted to know whether individual dynein trajectories displayed variations in the number of discrete instances (visits) negative α was observed during a single trajectory, and found this comparison to also remain nearly constant throughout different conditions (Supplemental Table S1). Combined, we rejected possibility 1 as a source of variation in negative α. We also rejected possibility 3, because we observed variation in prophase and anaphase negative α spectrum in the anterior and posterior that is consistent with force generation asymmetry in the zygote in previous studies (Supplemental Table S1, Rodriguez-Garcia 2018, Park 2008). Finally, we measured the average number of dynein particles that exhibited negative α behavior. Here, we found marked variation amongst conditions (Figure 5 D, Supplemental Table S1). These variations were consistent with the average time spent in negative α calculated in Figure 3E, suggesting variation in negative α spectra is a consequence of the number of dynein particles displaying an instance of negative α behavior. Importantly, there is a linear relationship between the number of dynein particles displaying negative α behavior and the total number of dynein particles in the imaging field. (Supplementary Table S1). Together, we find that the number of dynein particles able to bind to the cortex and display any negative α behavior (i.e. binding rate) is the aspect of dynein behavior that dictates the magnitude of cortical dynein force generation. Moreover, the number of dynein displaying negative α is positively correlated to the number of dynein particles at the cortex, suggesting a mechanosensitive cue that can tune force generation by recruiting a greater number of dynein particles to the cortex.

## Discussion

Here, we have utilized high spatiotemporal resolution fluorescence imaging with single particle tracking and a wMSD analysis to tease apart the functional relationship between dynein behavior and cortical force generation during pronuclear migration and anaphase. We have been able to show in distinct ways that a negative α corresponds to cortical force generation (Figure 3A, Figure 4B). The time a dynein particle spends displaying negative α is about two-fold greater in the anterior compared to the posterior segment of the zygote, consistent with the asymmetric force generation measurements observed by others during migration in the *C. elegans* zygote (Figure 3A) (Rodriguez-Garcia *et al*., 2018). In addition, depleting a subunit of the dynein complex (DLC-1), a dynein cofactor (dynactin), or the anchoring protein that links dynein to the force generation complex (LIN-5) causes a decrease in negative α behavior in the anterior (Figure 3B, 3C).

Although we have previously shown that both LIS-1 and SUR-6 depletions decrease cortical dynein motoring capacity (force generation), LIS-1 and SUR-6 depletions increase the amount of time dynein particles display negative α (Boudreau *et al*., 2019). We find that the increase of cortically tethered dynein in *lis-1* (RNAi) and *sur-6* (RNAi) zygotes is a result of an increase in dynein recruitment to the cortex (# of dynein particles per embryo: control (L4440) = 149.58, *lis-1* (RNAi) =203.53, *sur-6* (RNAi) = 271.94).

LIS-1 has been shown to positively regulate cytoplasmic dynein motoring capacity by modulating the ATPase activity of its AAA +domain (ATPases are associated with diverse cellular functions), causing dynein to be more stably bound to the microtubule for a longer period of time (Huang *et al*., 2012). In the absence of LIS-1, the dynein binding and unbinding rate to microtubules is increased. From the perspective of cortical dynein, the absence of LIS-1 would induce fast unbinding from microtubules, such that we observe greater time spent in negative α (tethered to the cortex), but dynein dissociates from the microtubule before it is capable of generating force.

PP2A-B55/SUR-6 is an important serine/threonine mitotic phosphatase. While we have evidence that SUR-6 does not interact with dynein directly, LIN-5 is a strong candidate for interaction with SUR-6 (Boudreau *et al*., 2019). Multisite phosphorylation of LIN-5 has been shown to regulate spindle position and chromosome regulation, and there is mounting evidence that an interplay between mitotic kinase and phosphatase turnover is the mechanism that controls this process (Portegijs *et al*., 2016, Keshri *et al*., 2020). This would mean that the phosphorylation state of LIN-5 allows it to bind and unbind to GPR 1/2, or to bind and unbind to dynein to generate force. Given that CDK-1 phosphorylates LIN-5 at T181, which is within the dynein binding domain of LIN-5, PP2A-B55/SUR-6 likely dephosphorylates this same locus, directly affecting the interaction between dynein and LIN-5 (Portegijs *et al*., 2016, Kotak *et al*., 2013). These considerations would explain the greater average time spent negative observed in *sur-6* (RNAi) zygotes. LIN-5 would be able to bind to GPR 1/2, but its interaction with dynein would be perturbed. Thus, we see dynein bound to the cortex and observe negative α behavior without generating any force on microtubules. Additional work is needed to confirm the locus within a LIN-5 binding domain that is dephosphorylated by SUR-6 to regulate dynein mediated force generation. Finally, our statistical analysis showed that cortical dynein behavior was aberrated in the anterior. The range of the distribution of dynein behavior became stochastic upon depleting SUR-6, indicating that dynein interaction with the cortex is not regulated in the depletion (Figure S2D). Our statistical analysis did not identify the same deregulation in the posterior, where microtubules emanating from the centrosomes are very dense in comparison to the anterior. This provides further evidence that SUR-6 directly interacts with the cortical force generation complex as a cell cycle regulator of force generation, since force generation primarily occurs in the anterior during prophase.

Our data indicates that in the *C. elegans embryo*, dynein recruitment to the cortex is modified in response to perturbation of proteins that are responsible for dynein mediated cortical force generation. When cortical pulling forces are dampened, recruitment of dynein to the cortex is increased, although depleting DLC-1 does not elicit the same response. We imagine a scenario where depletion to dynein regulators uncouples the actomyosin and microtubule cytoskeletal networks, such that depletions to regulatory proteins of dynein cortical force generation cause a higher posterior localization patterning, as seen in LIN-5, LIS-1 and SUR-6 depleted zygotes (Figure 3). We hypothesize these depletions abrogate the polarity of the microtubule cytoskeleton, while the actomyosin network is unaffected. The process could be a result of a membrane tension-sensing pathway that increases cortical dynein residence. Future work will be needed to address the mechanistic link between cortical dynein localization, residency, and cortical force generation production.

We report that there is less cortical dynein population in *dnc-1* (RNAi) than in *dlc-1* (RNAi). During embryogenesis, actomyosin cortical flows normally lead to a polarized accumulation of dynein at the cortex. We expect that *dnc-1* and *dlc-1* RNAi depletions do not affect actomyosin cortical flows. Thus, the difference in cortical dynein accumulation may be due to an auxiliary microtubule-based mechanism. Dynein may penetrate the cortical meshwork from the adjacent cytoplasm by interacting with microtubule associated plus tip proteins. Therefore, the number of dynein present in the meshwork in *dlc-1* (RNAi) embryos is similar to control (L4440) because dynein’s ability to interact with the microtubule plus end is not affected by the depletion. In contrast, in the DNC-1 depletion, dynein is unable to utilize the cortical microtubule meshwork for dynein localization and is unable to to interact with plus tip complexes (Markus and Lee 2011).

In summary, we have characterized the behavior of cortical dynein by combining high spatiotemporal fluorescence microscopy with a wMSD analysis. We have shown that negative α behavior in the x, y plane and negative α in fluorescence intensity provides an approximation for the behavior response for dynein’s ability to generate force, leading us to propose a simple spring-like model for understanding cortical dynein force generation. We also offer a mechanistic role for PP2a-B55/SUR-6 in regulating cortical force generation through the dephosphorylation of LIN-5. Further studies will be necessary to confirm SUR-6 dephosphorylation of LIN-5. Finally, we have evidence to suggest the existence of a mechanosensitive pathway that modifies cortical dynein localization in response to changes in cortical force generation.

## Materials and Methods

### C. elegans use, RNAi, and microscopy

The worm strain SV1803 (he264[eGFP::dhc-1]) was grown and maintained at 20°C using standard procedures. Bacterial strains containing a vector expressing double-stranded RNA under the isopropyl β-D-1-thiogalactopyranoside promoter were obtained from the Ahringer library (Bob Goldstein’s laboratory, University of North Carolina at Chapel Hill, Chapel Hill, NC). Targets were confirmed by sequencing.

Embryos were mounted in egg buffer between a 1.5 coverslip and a 4.5% agarose pad in egg buffer and a microscope slide and sealed with Valap. Embryos were imaged on a Nikon TIRF microscope with a 1.5x magnifier using a 100× Apo TIRF oil-immersion objective (Nikon), an Andor iXon3 EMCCD camera and NIS-Elements (Nikon) at 22°C.

### Image Analysis and Diffusion Coefficient Calculations

All image analysis was done using the TrackMate (Tinevez et al., 2017) plugin in FIJI (Schlinden et al., 2012). Tracking was performed using a Difference of Gaussian (DoG) detection segmenter and simple Linear Assignment Problem (LAP) tracking. Image processing was performed using a custom FIJI plug-in that is available upon request. To determine sections of a dynein’s trajectory corresponding to a force generation event, a windowed mean square displacement was performed. This analysis technique is implemented using a sliding window approach. A sliding window selects a number of points in a trajectory and calculates the mean square displacement over those points. A diffusion coefficient is then calculated for that section of the trajectory covered by the window. The window is then moved one point forward in time along the length of the trajectory.

### Mean Square Displacement

A mean square displacement (MSD) is performed on each windowed section of a given dynein’s trajectory. The MSD calculates the average distance travelled over various time lags, so that the average displacement every ***n***^th^ time step (where ***n*** is a whole number), and so on can be determined. MSD takes the form,

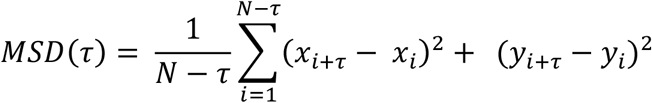

where N is the total number of points in the trajectory, **τ** is the time lag, (*x*_*i*,_ *y*_*i*_) is the position at time i, and (*x*_*i*+*τ*_, *y*_*i*+*τ*_) is the position i+**τ** units of time away.

### Windowing

A sliding window approach was used to calculate the MSD and diffusion coefficient throughout a given dynein’s trajectory. A window size of 6, corresponding to approximately 0.162 seconds, was used in this study. The length of each trajectory was extended by 10 points, where five copies of the initial position were concatenated to the beginning of the trajectory, and five copies of the final position were concatenated to the end of the trajectory.

### wMSD analysis

The motor protein dynein consumes energy through ATP hydrolysis and thus is able to move itself, as well as interact with the microtubule cytoskeleton and cell cortex. We reason that a sufficiently long trajectory includes dynein engaging in a number of these activities. Because of this we consider trajectories to consist of two general parts, an active part that reflects cortical interactions and motoring, and a passive component corresponding to dynein dwelling. The active and passive components of a trajectory manifest in displacement length, that the displacements between positions reflects dynein’s activity at the cortex. Axonemal dynein motors along a microtubule at a rate of 14 microns/second *in vitro*, which would result in displacements of 378nm for a sampling rate of 0.027 seconds. In contrast, to engage in force generation dynein is bound to the cortex and a microtubule, essentially confined to a single location during this process. These trajectories are most similar to Lévy flights, we expect that a trajectory will consist of both small and large displacements. Because the displacement measured from the step size of dynein motoring along a microtubule is so much greater than the displacement when dynein is engaged in force generation, we use a power law distribution to model the dynein trajectories’ MSD using a Lévy flight. For the mean square displacement, the Lévy flight takes the form

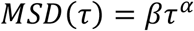

*β* is a generalized diffusion constant, and *α* quantifies the super-diffusion. The term super-diffusion is used since long steps are more likely than the probability given by a Brownian model. We interpret *β* as being descriptive of the characteristics of the media dynein is moving through, and *α* as characterizing the step length of dynein. *α* is related to the probability of a step length 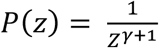 where z is the step length and, for instance, *α* = 3 − *γ* when 1 ≤ *γ* < 2 (Barthelemy *et al*., 2008).

By analyzing the trajectories using the wMSD and power law model we expect to be able to pick out points in the trajectory where dynein is undergoing cortical force generation. When dynein is tethered to the cortex, we expect to see slight displacements about a fixed point in 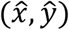 due to the length of the ‘tether’ and transient invaginations in the cell membrane. This would result in the dynein molecule’s fluorescent tag passing through the same location repeatedly in a manner similar to the motion of a spring. Since dynein is tethered to the cortex and passing through the same location, we expect to see subdiffusion, indicated by a negative *α* value. Subdiffusion has been shown to result from errors in the location of 30-nm gold colloids fixed onto a substrate, an experimental system similar to dynein tethered to the cortex, where errors in position were on the order of hundreds of nanometers (Martin *et al*., 2002). Dynein cortical force generation requires the molecule to be tethered to the cortex, and can be identified within a trajectory using the wMSD and looking for *α* < 0. To measure *α* we take the natural log of the power law to get, *log* (*MSD*(*τ*)) = *log* (*β*) + *αlog* (*τ*) and calculate *α* using a least square fit of a polynomial degree 1.

The use of windowing results in a spectrum of *α* values describing overlapping sections of the trajectory. For a trajectory of length N, there are N+ 10 points added for a window size of 6 (N + 2(window length – 1)), resulting in an *α* spectrum containing N+5 values (N + (window length-1)). To determine the amount of time a dynein exhibits subdiffusive behavior (*α* < 0) find the difference in the index *i* position in the *α* spectrum of each *α* < 0 in order and compute,

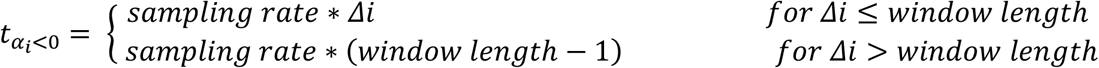

for each, where Δ*i* is the difference in index position within the spectra between successive negative values. Then sum for all 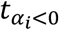, so that the total time spent subdiffusive for a given dynein trajectory is 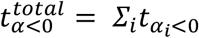. To calculate the total time every dynein spends subdiffusive in a side of the cell 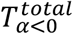, calculate

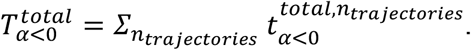

Where *n*_*trajectories*_ is the number of dynein trajectories collected in a single movie of either anterior or posterior ends of the cell. To characterize a given depletion, take the average of 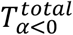 for each replicate in that group to get 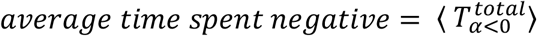.

### Bootstrapping and Two-Sample Kolmogorov-Smirnov

To determine the likelihood of measuring the average time spent negative (ATSN) by chance, we performed a bootstrapping analysis. Each trajectory from every replicate had its position coordinates shuffled 20 times using a random number generator in MatLab. These shuffled trajectories were then analyzed using the same method as the experimental data presented in this paper, such that there are 20 replicates per movie. The average value of each replicate for every movie was then taken and the distribution analyzed using a box plot to compare it with the data measured.

The distributions were compared using a two sample Kolmogorov-Smirnov test to determine if the bootstrapped values and the experimental values come from the same distribution. The null hypothesis for the two-sample Kolmogorov-Smirnov test is that the two sets of data have the same distribution. It then returns either a “0” if it was unable to reject the null hypothesis or a “1” if the test rejects the null hypothesis at the 5% significance level.

### Numerical Programming

All numerical programming was performed using Matlab version R2018b.

## Acknowledgements

We greatly thank Jennifer Heppert (Goldstein lab, UNC–Chapel Hill) for reagents, Tony Perdue (Microscopy Core, Biology Department, UNC–Chapel Hill) for support. We also thank members of the Amy Maddox lab (UNC—Chapel Hill) and Paul Maddox lab (UNC—Chapel Hill), as well as Sebastian Fürthauer (Flatiron Institute, Center for Computational Biology) for critical reading and discussion of the manuscript. This study was supported by the National Science Foundation CAREER Award 1652512 to P.S.M. J.B.L. was supported in part by a grant from NIGMS under award T32 GM119999, as well as NIH R01-102390 and NSF 1616661.

## Author contributions

A.E. and V.B designed and carried out experiments in *C. elegans* embryos and performed image and data analysis. J.B.L. wrote MatLab based code used for data calculations, statistics, and computational modeling, and contributed to data analysis. A.E, J.B.L. and P.S.M. wrote the article.

