## Supplemental Figure 1 for "Single-particle tracking of dynein identifies PP2A B55/SUR-6 as a cell cycle regulator of cortical force generation"

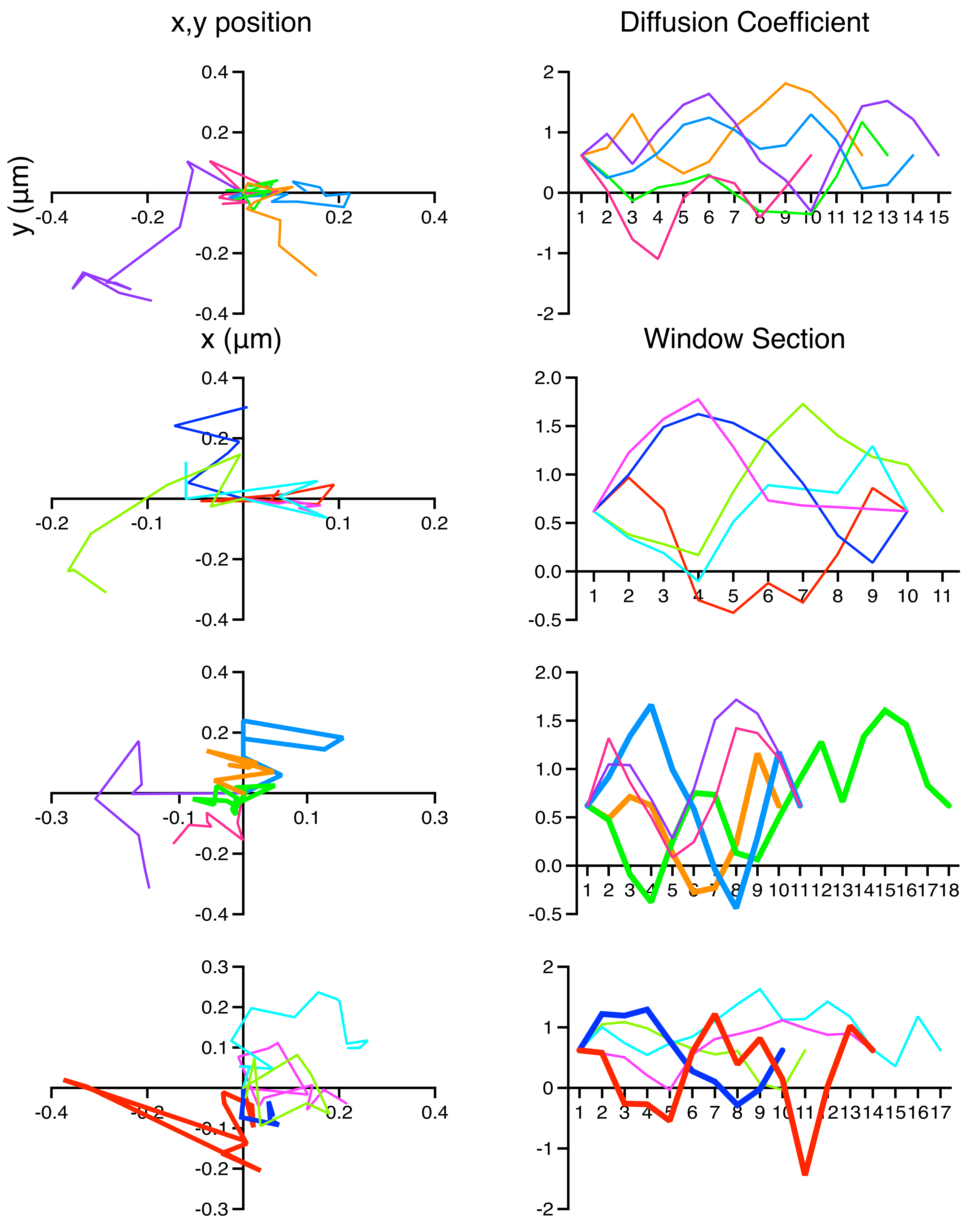


Supplemental Figure 1. A selection of twenty randomly chosen diffusion tracks. x,y position of 20 random trajectories with extrapolated diffusion coeffiient spectra using a wMSD analysis (split in to 4 panels for simplicity). In the bottom two panels, tracks with α < 0 are highlighted to compare tracks with multiple negative α values to those without.
