## Supplemental Figure 2 for "Single-particle tracking of dynein identifies PP2A B55/SUR-6 as a cell cycle regulator of cortical force generation"

A

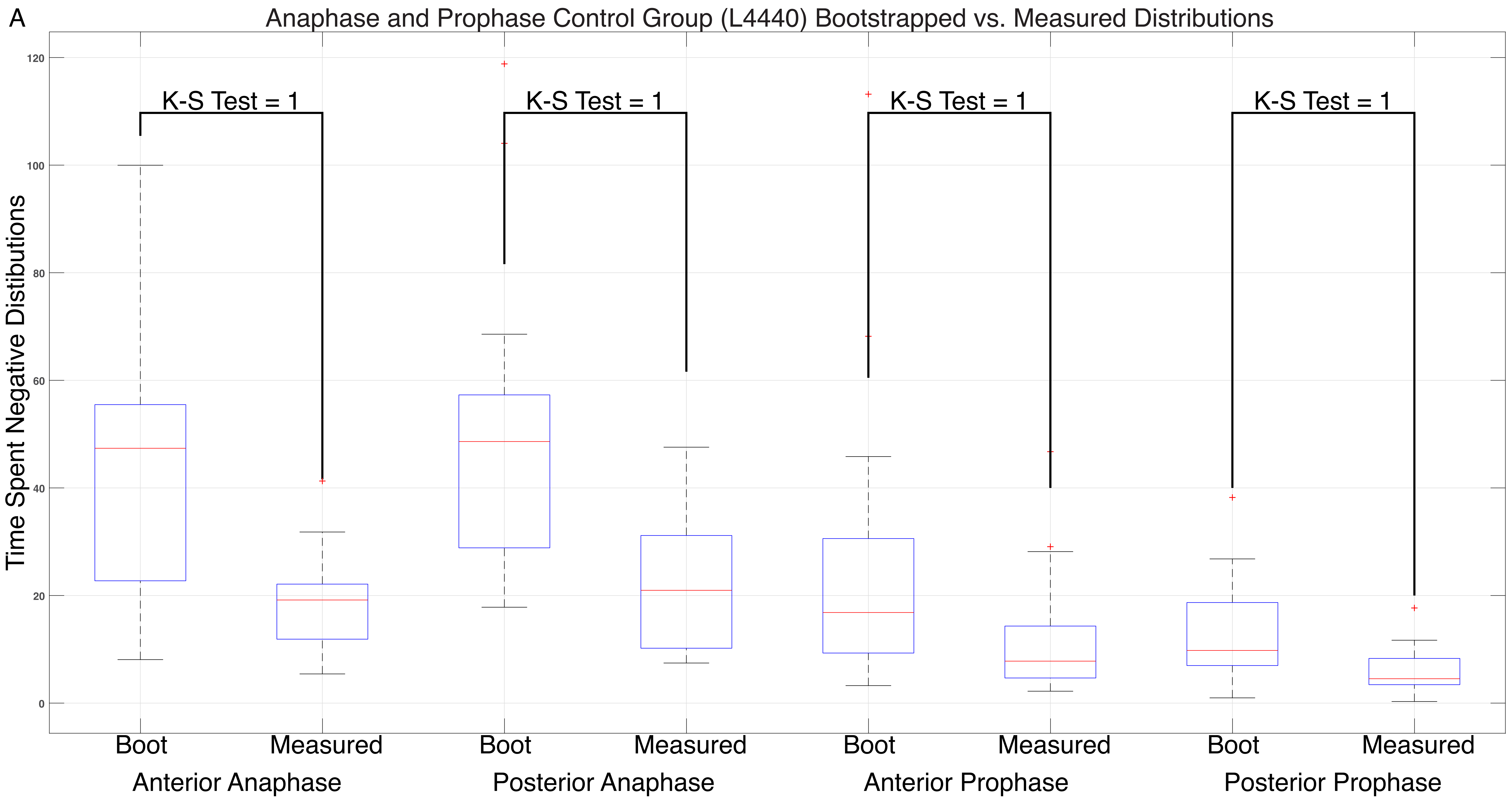

Anaphase and Prophase Control Group (L4440) Bootstrapped vs. Measured Distributions
Time Spent Negative Distibutions
120
100
80
60
40
20
0
K-S Test = 1
K-S Test = 1
K-S Test = 1
K-S Test = 1
Boot
Measured
Boot
Measured
Boot
Measured
Boot
Measured
Anterior Anaphase
Posterior Anaphase
Anterior Prophase
Posterior Prophase

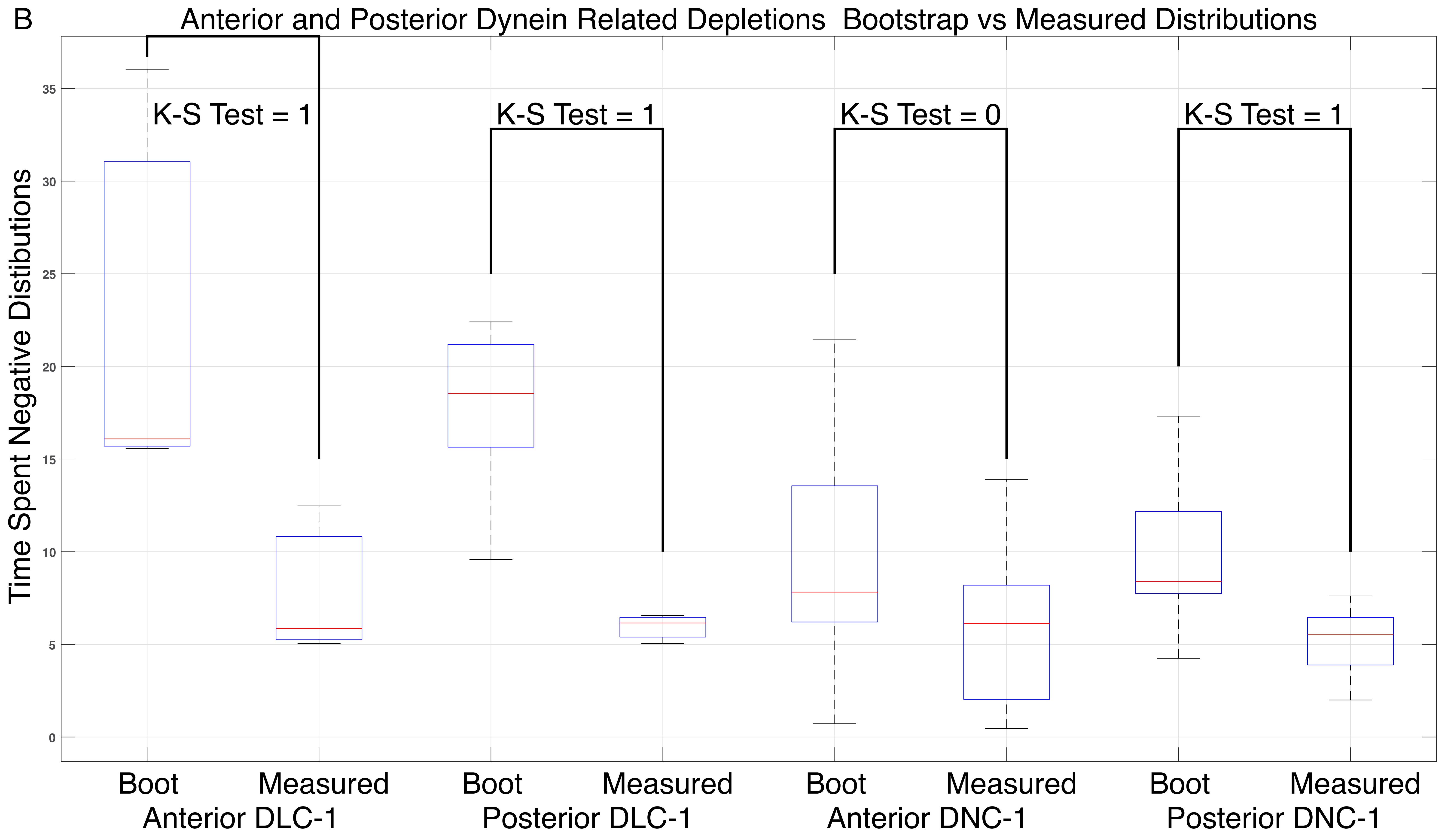

B
Anterior and Posterior Dynein Related Depletions Bootstrap vs Measured Distributions
Time Spent Negative Distibutions
K-S Test = 1
K-S Test = 1
K-S Test = 0
K-S Test = 1
0
5
10
15
20
25
30
35
Boot
Measured
Anterior DLC-1
Boot
Measured
Posterior DLC-1
Boot
Measured
Anterior DNC-1
Boot
Measured
Posterior DNC-1

C

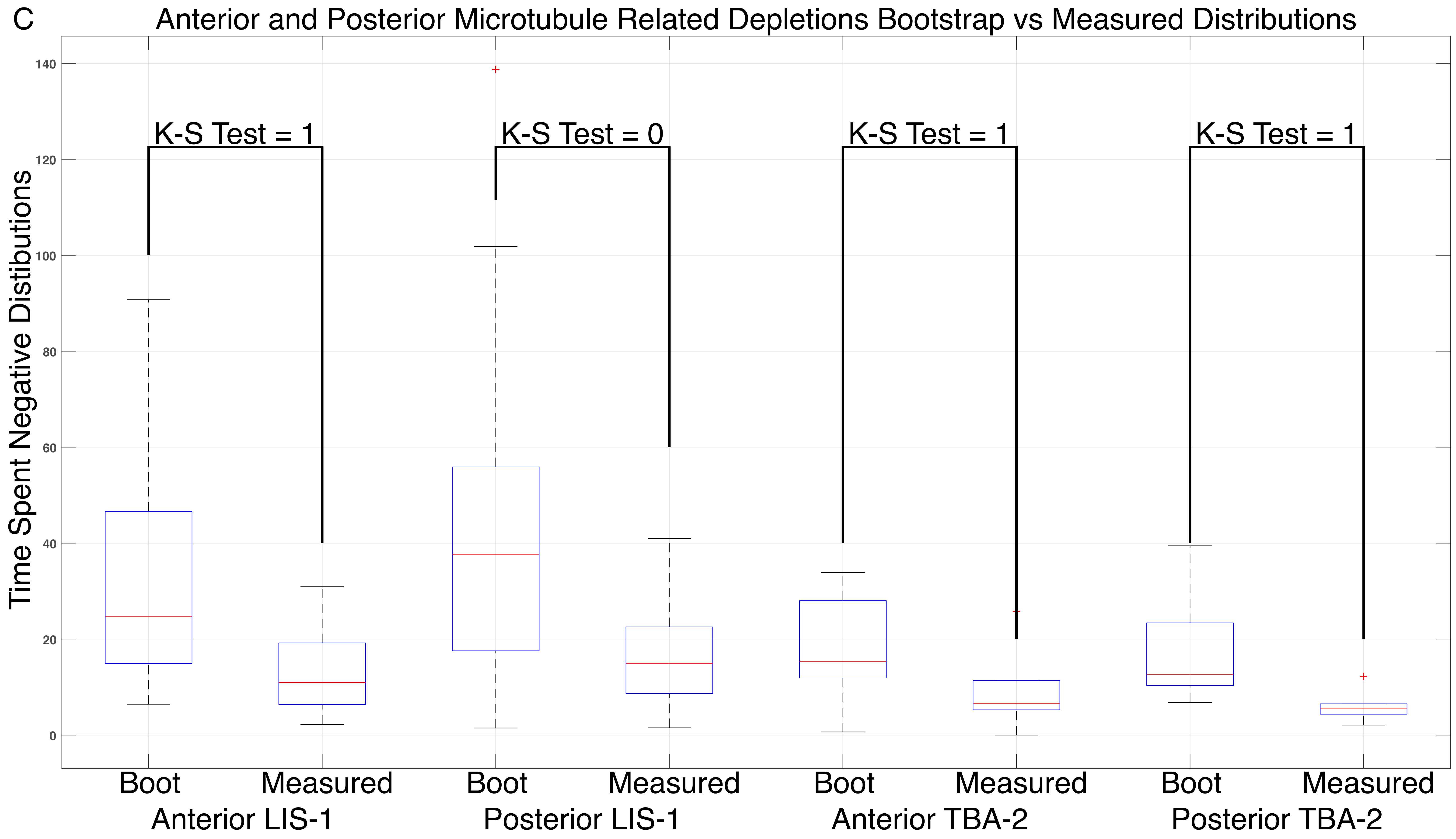

Anterior and Posterior Microtubule Related Depletions Bootstrap vs Measured Distributions
Time Spent Negative Distibutions
140
120
100
80
60
40
20
0
K-S Test = 1
K-S Test = 0
K-S Test = 1
K-S Test = 1
Boot
Measured
Boot
Measured
Boot
Measured
Boot
Measured
Anterior LIS-1
Posterior LIS-1
Anterior TBA-2
Posterior TBA-2

D

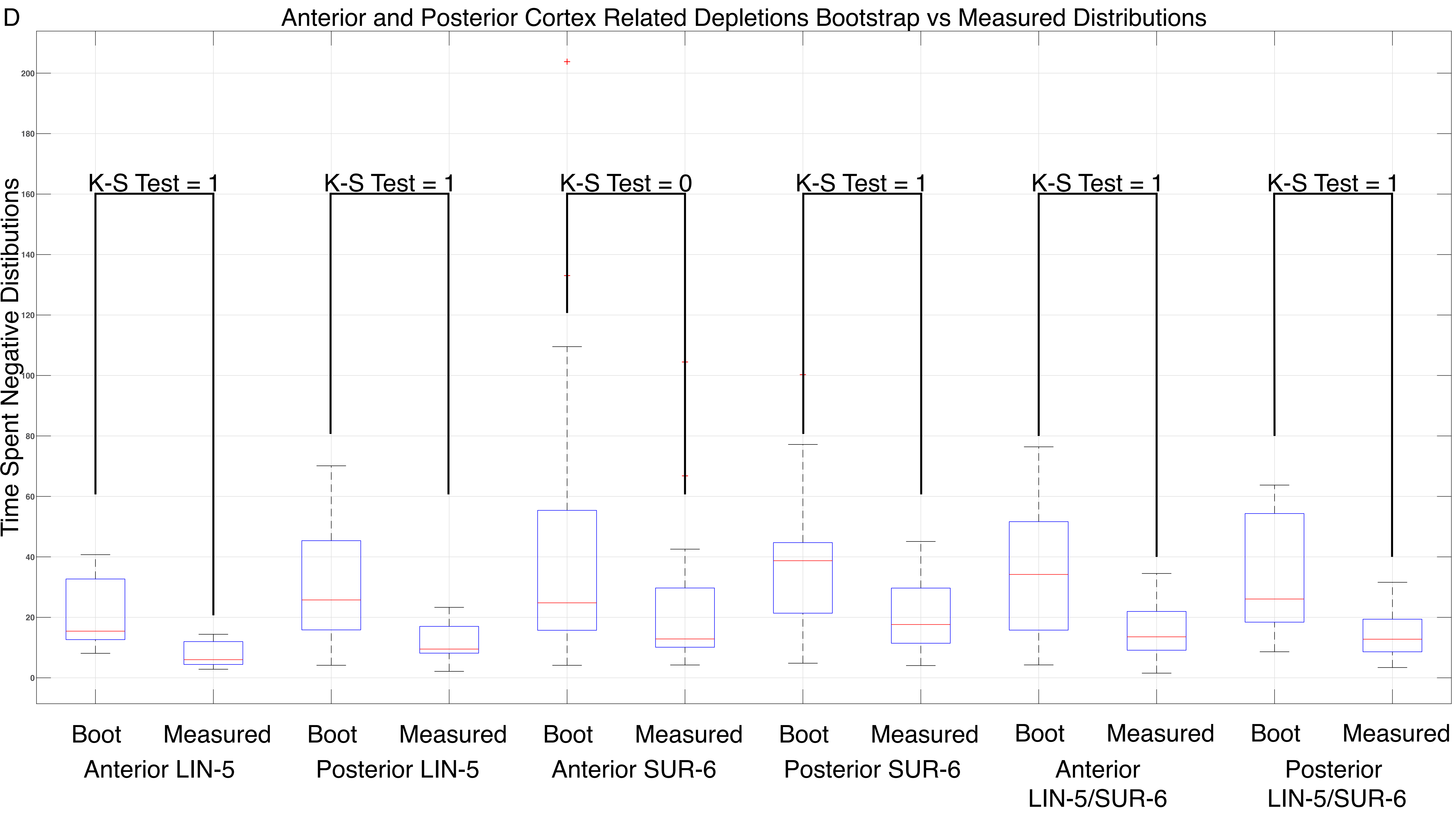


Supplemental Figure 2D. A-D: Bootstrap statistical analysis. K-S Test = 1 rejects the null hypothesis (see materials and methods).
