## Supplemental Table 1 for "Single-particle tracking of dynein identifies PP2A B55/SUR-6 as a cell cycle regulator of cortical force generation"

| | Mean Time Tethered | Median Time Tethered | Visits (mean) | Fraction of Trajectories that are at any point cortically tethered | # Dynein engaging in Force Generation/replicate number | Total Dynein per replicate number | Length ≥ 6 (Total Trajectories Analyzed) | Length ≥ 10 | Length ≥ 15 |
|---|---|---|---|---|---|---|---|---|---|
| **Anterior Anaphase** | 0.1876 | 0.189 | 1.35 | 0.3441 | 91.2308 | 265.1538 | 2475 | 831 | 269 |
| **Anterior Prophase** | 0.1895 | 0.189 | 1.3788 | 0.3838 | 57.4167 | 149.5833 | 2552 | 853 | 272 |
| **Posterior Anaphase** | 0.1879 | 0.189 | 1.3561 | 0.3642 | 104.7143 | 287.5 | 2934 | 1003 | 349 |
| **Posterior Prophase** | 0.187 | 0.189 | 1.2525 | 0.3237 | 30.3043 | 93.6087 | 1450 | 335 | 81 |
| **Anterior** | | | | | | | | | |
| **DLC-1** | 0.1848 | 0.162 | 1.2047 | 0.2893 | 42.33 | 146.33 | 303 | 96 | 27 |
| **DNC-1** | 0.1909 | 0.189 | 1.4103 | 0.3772 | 27.8571 | 73.8571 | 326 | 70 | 14 |
| **TBA-2** | 0.1866 | 0.189 | 1.4198 | 0.2213 | 41.5556 | 187.7778 | 681 | 223 | 79 |
| **LIS-1** | 0.1879 | 0.189 | 1.3199 | 0.3274 | 66.6316 | 203.5263 | 2723 | 819 | 258 |
| **LIN-5** | 0.1871 | 0.189 | 1.2518 | 0.2931 | 41.8462 | 142.7692 | 1265 | 321 | 90 |
| **SUR-6** | 0.1905 | 0.189 | 1.4241 | 0.4203 | 114.2941 | 271.9412 | 3419 | 1241 | 431 |
| **SUR-6/LIN-5** | 0.1888 | 0.189 | 1.3743 | 0.3794 | 78.4286 | 206.7143 | 2113 | 684 | 233 |
| **Posterior** | | | | | | | | | |
| **DLC-1** | 0.1804 | 0.162 | 1.1953 | 0.2826 | 33.8 | 119.6 | 418 | 113 | 24 |
| **DNC-1** | 0.1872 | 0.189 | 1.4257 | 0.3156 | 24.6667 | 78.1667 | 287 | 54 | 14 |
| **TBA-2** | 0.1809 | 0.162 | 1.2732 | 0.1764 | 32.3333 | 183.3333 | 1450 | 335 | 81 |
| **LIN-5** | 0.1856 | 0.162 | 1.2589 | 0.3175 | 60.3077 | 189.9231 | 3020 | 944 | 307 |
| **LIS-1** | 0.187 | 0.189 | 1.317 | 0.3297 | 91.0667 | 276.2 | 1732 | 510 | 153 |
| **SUR-6** | 0.1933 | 0.189 | 1.5016 | 0.4184 | 91.7059 | 219.1765 | 2754 | 1015 | 354 |
| **SUR-6/LIN-5** | 0.1869 | 0.189 | 1.361 | 0.3663 | 71.5294 | 195.2941 | 2460 | 795 | 281 |
